# Trim33 conditions the lifespan of primitive macrophages and onset of definitive macrophage production

**DOI:** 10.1101/2022.04.06.487382

**Authors:** Doris Lou Demy, Anne-Lou Touret, Mylène Lancino, Muriel Tauzin, Lavinia Capuana, Constance Pierre, Philippe Herbomel

## Abstract

TRIM33 (Tif1-γ) is a transcriptional regulator notably involved in several aspects of hematopoiesis. It is essential for the production of erythrocytes in zebrafish, and for the proper functionning and aging of hematopoietic stem and progenitor cells (HSPCs) in mice. Here we have found that in zebrafish development, Trim33 is essential cell-autonomously for the lifespan of the yolk sac derived primitive macrophages, as well as for the initial production of definitive (HSPC-derived) macrophages in the first niche of definitive hematopoiesis, the caudal hematopoietic tissue. Moreover, Trim33 deficiency leads to an excess production of definitive neutrophils and thrombocytes. Our data indicate that Trim33 radically conditions the differentiation ouput of aorta-derived HSPCs in all four erythro-myeloid cell types, in a niche-specific manner.

## INTRODUCTION

Wandering within most tissues from early on in ontogeny, tissue-resident macrophages are often pictured as sentinels, always on alert, with the mission to continually detect and adequately react to any fluctuations within the surrounding tissues, from the small and natural consequences of proper organ functioning to extreme pathological/inflammatory aggressions. These roles require a high level of transcriptional versatility, to provide macrophages with a wide range of possible reactions and quick reactivity. As it is thought that, at least in mammals, tissue-resident macrophages populations maintain themselves throughout life, they also need robust self-renewal capacities and a long lifespan. The transcriptional toolbox necessary for macrophages to face so diverse tasks for such an extended period of time is still poorly known.

Macrophages arise in successive waves in vertebrate ontogeny – first in the yolk sac, and later within the developing larva or foetus as one of the differentiation outcomes of the hematopoietic stem/progenitor cells (HSPCs), in the successive niches where the so-called “definitive” hematopoiesis takes place. In zebrafish development, the initial – or “primitive” – wave of macrophages originates from the anterior-most lateral mesoderm. By mid-somitogenesis, myeloid progenitors arise directly from this mesoderm and migrate into the adjacent yolk sac to differentiate into primitive macrophages and neutrophil precursors, which then quickly invade the embryo’s mesenchyme and then the epidermis. Only the macrophages further colonize the brain and retina, to become the primitive microglia (Herbomel et al., 1999; Herbomel et al., 2001; Le Guyader et al., 2008).

Definitive hematopoiesis begins in the embryo with the emergence of HSPCs from the endothelium of the ventral wall of the aorta in the trunk (Kissa & Herbomel, 2010). These HSPCs then home to a first niche in the tail, the Caudal Hematopoietic Tissue (CHT) where they expand and undergo multi-lineage differentiation, including in lymphoid cells, and later home to the kidney, the niche of lifelong hematopoiesis (Murayama et al., 2006; Kissa et al., 2008). The CHT then the “kidney marrow” thus are the hematopoietic niche homologs in fish of the fetal liver then bone marrow in mammals.

The transcription co-factor Trim33/Tif1-γ has been known for years to be an important actor of hematopoiesis. It regulates transcription by binding to various DNA-binding transcription factors, such as SMADs, PU.1 and Scl/Tal1 (Ferri et al., 2015; Kusy et al., 2011; Xi et al., 2011). In zebrafish, it was first identified as essential cell-autonomously for the differentiation of the primitive erythrocytes from the embryo’s mesoderm (Ransom et al., 2004). *Moonshine* mutant embryos, which harbor a null mutation in Tif1-γ/Trim33, appear “bloodless”, as the primitive erythroid progenitors undergo early apoptosis by mid-somitogenesis. Monteiro et al. (Monteiro et al., 2011) later found that the definitive, HSPC-derived erythropoiesis normally occurring in the CHT is also defective in *moonshine* swimming larvae, while granulopoiesis is enhanced. In mammals, the hematopoietic role of Trim33/Tif1-γ has only been studied in postnatal/adult mice. Trim33-deficient HSPCs showed signs of premature aging (Quéré et al., 2014) and an increased capacity to generate myeloid progenitors - to the expense of other lineages (Aucagne et al., 2011), that showed a diminished capacity to differentiate in macrophages (Chrétien et al., 2016; Gallouet et al., 2017).

More recently, Trim33 has been shown to be necessary for several essential functions of differentiated macrophages. In mice, Romeo and coworkers found that Trim33-deficient macrophages are unable to resolve inflammation (Ferri et al., 2015; Gallouet et al., 2017; Petit et al., 2019). In the zebrafish, we found that in *moonshine* embryos, primitive macrophages and neutrophils are produced, and then disperse throughout the embryo’s interstitium, but they display a reduced basal mobility, and are unable to respond to developmental or inflammatory recruitment signals. This notably leads to an absence of primitive microglia (Demy et al., 2017).

We now report that over the next two days of development of *moonshine* larvae, the primitive macrophages (but not the neutrophils) prematurely disappear. In addition, the first wave of definitive macrophage production from aorta-derived HSPCs is missing, while the production of HSPC-derived neutrophils and thrombocytes is boosted.

## RESULTS

### *Moonshine* primitive macrophages disappear prematurely between 54 hpf and 4 dpf

When proceeding with the characterization of the myeloid phenotype of our own *moonshine* (Trim33 null mutant) allele, *mon^NQ039^* (Demy et al., 2017), we discovered than following the strong navigation defect displayed by both primitive macrophages and neutrophils by 2-3 days post-fertilization (dpf), the macrophages (but not the neutrophils) had mostly disappeared altogether from the mutant larvae by 4 dpf (**Fig. 1A;** zebrafish embryos become “swimming larvae” after hatching, by 2.5 dpf). We confirmed this previously unreported phenotype of *moonshine* by counting mCherry positive (mCherry+) macrophages at 4 dpf in *mon^NQ039^ Tg(mpeg1:mCherry-F)* larvae, both manually on live individual larvae (**Fig. 1B**) and by fluorescence associated cell sorting (FACS) of pools of >50 mutant larvae and their siblings (**Fig. 1C.** We had previously shown that *moonshine* mutant embryos have a normal number of macrophages and neutrophils at 48 hours post-fertilization (hpf) (Demy et al., 2017). So we looked more closely at the kinetics of this macrophage disappearance by revealing the macrophage population of *mon^NQ039^* mutant embryos and their siblings by whole-mount *in situ* hybridization (WISH) for *csf1ra* from 2 to 4 dpf (**Fig. S1A**), and we found that the disappearance of the mutant macrophages was very gradual, starting around 54 hpf. We obtained the same results via *in vivo* imaging of mon^*TB222*^ *Tg(mfap4:mCherry-F)* larvae (**Fig. 1D**). Following up individual larvae over time and counting their macrophages at 2, 3 and 4 dpf confirmed the gradual disappearance of their macrophages (**Fig. 1E**).

**Fig. 1.**
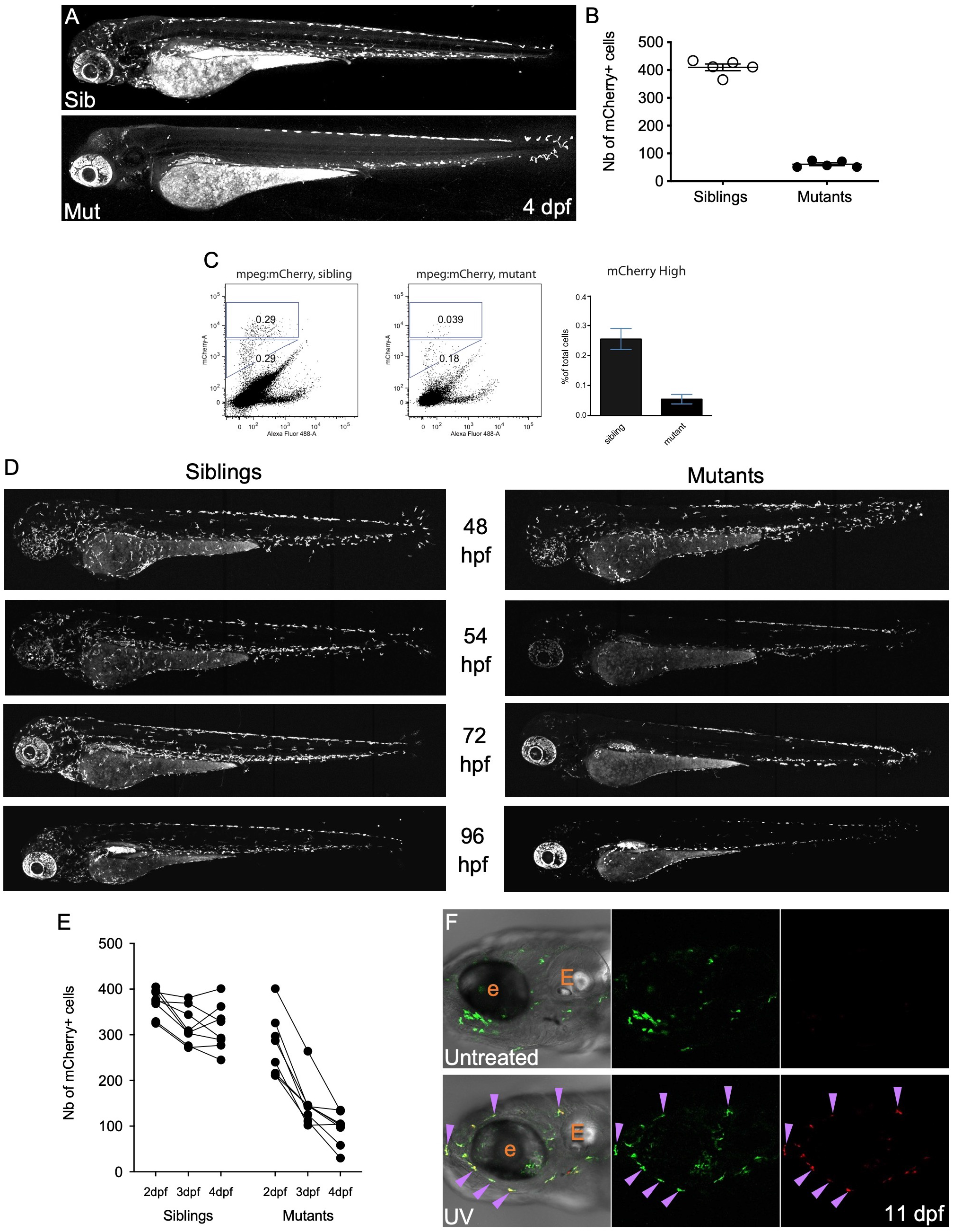
In *moonshine* mutant zebrafish, primitive macrophages prematurely disappear between 54 hpf and 4 dpf. **(A)** mCherry+ macrophages in live 4 dpf *Tg(mpeg1:mCherry-F)* control sibling (top) and *mon^NQ039^* mutant (bottom) larvae; maximum projection, lateral view. **(B)** Quantification of mCherry+ macrophages in live *Tg(mpeg1:mCherry-F)* sibling (white dots) and *mon^NQ039^* mutant (black dots) larvae at 4 dpf (n=5 larvae per condition). **(C**) Quantification of mCherry+ macrophages from *mon^NQ039^/mpeg1:mCherry-F* sibling (left plot) and mutant (right plot) larvae at 4 dpf by FACS. Plots: representative data; graph: pool of 2 experiments. **(D)** mCherry+ macrophages in live *Tg(mfap4:mCherry-F)* zebrafish larvae at 48, 54, 72 and 96 hpf in control siblings (left) and *mon^TB222^* mutants (right), maximum projection. **(E)** Quantification of total mCherry+ macrophages in single *mon^TB222^/Tg(mfap4:mCherry-F)* sibling and mutant larvae followed up from 2 to 4 dpf (n=8 larvae per condition). **(F)** Head region at 11 dpf of *Tg(mpeg1:Gal4;UAS:Kaede)* larvae - untreated (top) or UV-photoconverted at 2 dpf (bottom); single confocal planes. Purple arrowheads point at double positive (hence photoconverted) macrophages. e, eye; E, ear.

To check whether this disappearance of primitive macrophages in *moonshine* mutants is premature, we used Ultra-Violet (UV) exposure to completely photoconvert from green to red the Kaede proteins of *Tg(mpeg1:Gal4/UAS:Kaede)* embryos at 2 dpf. Upon UV-driven photoconversion all primitive macrophages become red (**Fig. S1B)**. By the next days, new (green) Kaede protein had been synthesized in these photoconverted macrophages, that now appeared yellow, as they contained both green and red Kaede protein (**Fig. S1B**). Numerous such cells were detected up to 9 days post photoconversion (**Fig. 1F**), meaning that the lifespan of wild-type (WT) primitive macrophages is at least 11 days, and that the disappearance of the whole primitive macrophage population that we witness in *moonshine* mutants between 3 and 4 dpf is indeed very premature.

### Primitive macrophages of Trim33 mutants accumulate morphological and metabolic defects before dying

Further *in vivo* observations and time-lapse imaging between 54 hpf and 4 dpf revealed that several cellular characteristics of *mon* primitive macrophages are lost upon their progressive disappearance. By 3 dpf, while most differentiated interstitial macrophages in WT larvae have acquired an elongated and ramified morphology, macrophages at similar locations in mutant larvae appear more round and less ramified (**Fig. 2A-C**). It seems that they also tend to loose adhesion capacity, as many are detected circulating in the heart and blood vessels, carried away by the blood flow (**Fig. 2D**). High-magnification *in vivo* observations of macrophages in *Tg(mfap4:mCherry-F) mon^TB222^* mutants and their siblings at 3 dpf also revealed a decrease of fluorescence in these cells prior to their disappearance (**Fig. 2E**), both in cell area (**Fig. 2F**) and total intensity (**Fig. 2G**). Quantification of the fluorescence of *mon^TB222^* macrophages expressing both mfap4:mCherry-F (membrane-adressed) and mfap4:turquoise (whole-cell reporter) transgenes *in vivo* (**Fig. 2H-I; Fig. S2A-B**) confirmed the global decrease of mCherry-F fluorescence in mutant macrophages, but also highlighted that this farnesylated fluorescent protein had mostly disappeared from the plasma membrane where it should be addressed (**Fig. 2I, black arrowheads**), and only persisted at the center of the cell (**Fig. 2I, purple arrowhead**). These mCherry+ hot spots within the mutant macrophages colocalized with numerous vesicles that were highly refractile upon Nomarski imaging (**Fig. 2J**), and acidic (**Fig. 2K**), suggesting that they were of phagocytic origin, unlike most of the mCherry+ intracellular hotspots of macrophages in sibling larvae **(Fig. 2K**). *In vivo* & time-lapse imaging of macrophages in *moonshine^TB222^ Tg(mfap4:turquoise)* mutants between 3 and 4 dpf then allowed us to witness dead cells still expressing macrophage driven fluorescence (**Fig. 2L**), and macrophages dying by bursting into several turquoise+ smaller pieces reminiscent of apoptotic bodies (**Fig. 2M**). Of note, macrophage death *in vivo* cannot be reliably assessed by the usual markers of apoptosis, for many of these professional phagocytes contain engulfed dead cell remnants that stain positive for these markers – hence our use of *in vivo* time-lapse imaging.

**Fig. 2.**
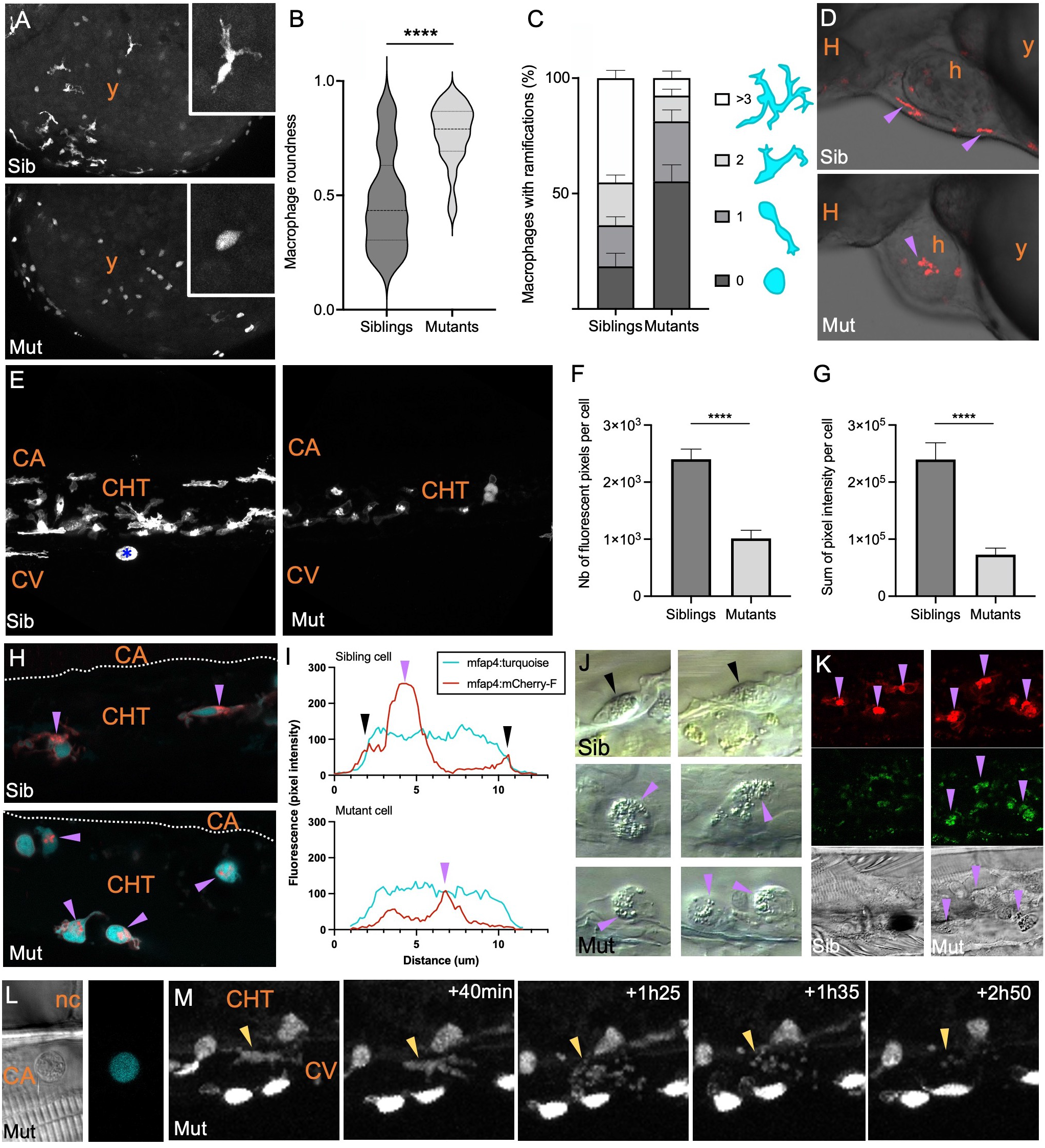
Primitive Trim33-deficient macrophages accumulate cytomorphological and metabolic defects before disappearing. **(A)** Turquoise+ macrophages in the yolk sac of live *mon^TB222^/Tg(mfap4:turquoise)* control sibling (top) and mutant (bottom) at 3 dpf; maximum projection. **(B)** Quantification of the roundness of turquoise+ macrophages in live *mon^TB222^*/*Tg(mfap4:turquoise)* sibling (left plot, dark grey) and mutant (right plot, light grey) larvae at 3 dpf (n=137 cells from siblings and 79 cells from mutants). **(C**) Percentage of turquoise+ macrophages with 0, 1, 2 or ≥ 3 ramifications in live *mon^TB222^/Tg(mfap4:turquoise)* sibling and mutant at 3 dpf (n=127 cells from siblings and 86 cells from mutants). **(D)** mCherry+ macrophages in the heart region of 3 days-old live *mon^TB222^/Tg(mfap4:mCherry-F)* control sibling (top) and mutant (bottom); single confocal plane, rostral to the left. Purple arrowheads point at macrophages. **(E)** mCherry+ macrophages in the caudal region of 3 days-old live *mon^TB222^/Tg(mfap4:mCherry-F)* control sibling (left) and mutant (right), maximum projection, rostral to the left. Blue asterisk labels a pigment cell. **(F-G)** Quantification of macrophage fluorescence in live *mon^TB222^/Tg(mfap4:mCherry-F)* sibling and mutant larvae at 3 dpf, based on **(G)** fluorescence area (nb of fluorescent pixels per macrophage) and **(F)** total fluorescence (sum of pixel intensities per macrophage); n=10 cells per condition. **(H)** Macrophages in the caudal region of 3 days-old live control siblings (top) and *mon^TB222^* mutants (bottom) expressing the *mfap4:turquoise* (macrophages, cyan channel) and *mfap4:mCherry-F* (macrophage membranes, red channel) transgenes; single confocal plane, rostral to the left. Purple arrowheads point at mCherry-F accumulation inside macrophages. **(I)** Quantification of fluorescence (pixel intensity) along cross-sections of macrophages in *mfap4:turquoise/Tg(mfap4:mCherry-F)* live sibling (top) and mutant (bottom) at 3 dpf. Black arrowheads point at membrane-associated mCherry-F signal surrounding the whole-cell turquoise signal in sibling cells. Purple arrowheads point at mCherry-F intracellular accumulation. Images used for plotting and other examples are shown in **Fig. S2**. **(J)** VE-DIC/Nomarski imaging of macrophages in the caudal region of 3 days-old live WT (2 top images) and *mon^NQ039^* mutants (4 bottom images); rostral to the left. Purple arrowheads point at refractile vesicles accumulation inside mutant macrophages. **(K)** Macrophages in the caudal region of 3 days-old live *mon^TB222^ Tg(mfap4:mCherry-F)* (macrophages, red channel) stained with LysoID-green (acidic compartments, green channel), control siblings (left) and mutants (right), single plane, lateral view. Purple arrowheads point at the mCherry-F intracytoplasmic accumulations that appear to be also acidic (LysoID+) and to coincide with the refractile vesicles in mutant macrophages. **(L)** Bright-field and Turquoise fluorescence signal of a dead macrophage in the trunk region of a live *mon^TB222^/Tg(mfap4:turquoise)* mutant at 72 hpf, single confocal plane, rostral to the left. **(M)** Selected time points of an *in vivo* time-lapse confocal imaging sequence in the caudal region of a *mon^TB222^/Tg(mfap4:turquoise)* mutant at 72 hpf (rostral to the left). Yellow arrowheads point at a macrophage that dies by bursting into apoptotic bodies. y, yolk sac; H, head; h, heart; CA, caudal artery; CHT, caudal hematopoietic tissue; CV, caudal vein; nc, notochord; Nb, number; um, micrometers.

### *Moonshine* larvae eventually recover macrophages from delayed definitive hematopoiesis

Observation of *moonshine* larvae later in development revealed that macrophages progressively reappeared in the mutants starting around 6 dpf (**Fig. 3A,C**; **Fig. S3A**), and that a full macrophage population was usually restored by 8.5 dpf (**Fig. 3B**). These macrophages seemed able to invade most larval tissues, including the brain where they settled to become microglia (**Fig. 3D**). To assess whether this later wave of macrophage production similarly occurred in WT larvae, we performed several experiments aiming at removing all primitive macrophages from the embryos and monitoring the (re)appearance of these cells in *mon* mutants and their siblings (**Fig. S3B**; **Fig. 3E**). First of all, we used two anti-sense morpholinos (Mo), targeting the transcription factors Pu.1 and Irf8, respectively, that have been shown to suppress primitive macrophage production, but not definitive (HSPC-derived) macrophages (Rhodes et al., 2005; Yu et al., 2017). Pu.1 Mo prevents all primitive myeloid cells formation (both primitive macrophages and neutrophils), whereas the Irf8 Mo disrupts the balance between these two myeloid cell fates, so that all progenitors become neutrophils. Both morpholinos resulted in a complete absence of macrophages at 48 hpf (**Fig. S3B**). We also took advantage of two techniques already used successfully in zebrafish to kill all macrophages at 48 hpf: clodronate-filled liposomes injection in the blood flow (Bernut et al., 2014), and metronidazole (Mtz) treatment of *Tg(mpeg1:Gal4; UAS:nfsB-mCherry)* embryos, wherein macrophages specifically express a nitroreductase that converts Mtz into toxic metabolites (Davison et al., 2007; Palha et al., 2013). We then monitored macrophage reappearance over time in these different conditions of primitive macrophage depletion, both in *mon^NQ039^* mutants (**Fig. 3E, left graph**) and WT larvae (**Fig. 3E, right graph**). Macrophage reappearance in *moonshine* mutants was pretty homogeneous amongst conditions and consistent with our previous observations, starting by 6 dpf (**Fig. 3E**, left graph). In contrast, macrophage reappearance occurred much earlier in WT larvae, placing the beginning of larval (definitive) macrophage production at 4 dpf (**Fig. 3E**, right graph).

**Fig. 3.**
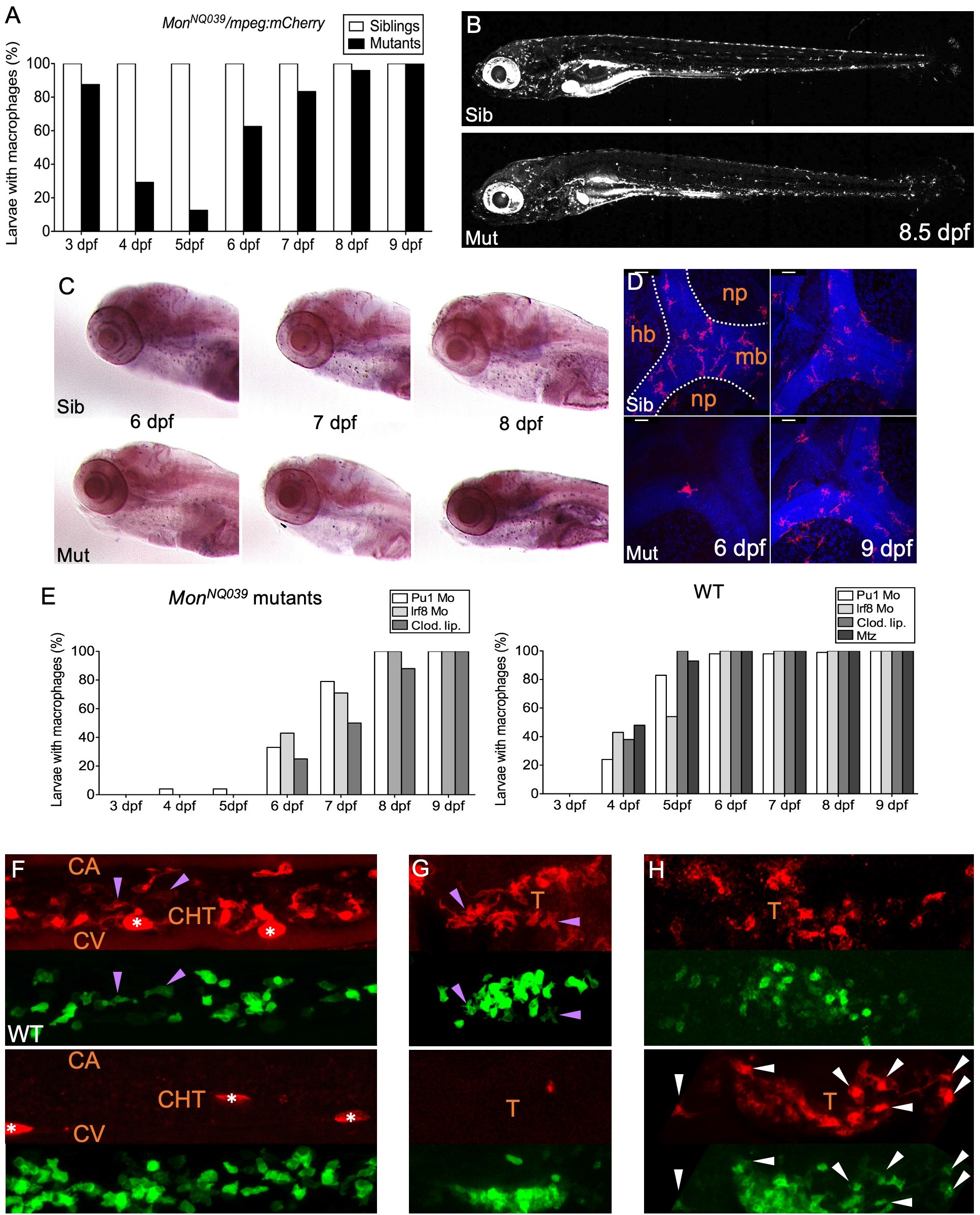
The production of macrophages from definitive hematopoiesis is delayed in *moonshine* mutants. **(A)** *In vivo* quantification of the percentage of sibling and *mon^NQ039^* mutant larvae with macrophages from 3 to 9 dpf, using a *Tg(mpeg1:mCherry-F)* background; n=84 sibling and 24 mutant larvae. **(B)** mCherry+ macrophages in live 8.5 dpf *Tg(mfap4:mCherry-F)* control sibling (top) and *mon^TB222^* mutant (bottom) larvae; maximum projection. **(C)** *In situ* hybridization for whole-mount detection of *Csfr1a*-expressing cells in control siblings (top) and *mon^NQ039^* mutants (bottom) at 6, 7 and 8 dpf. **(D)** Fluorescent immunodetection of L-plastin+ (leucocytes, red channel) macrophages in the brain of control siblings (top) and *mon^NQ039^* mutants (bottom) at 6 (left panels) and 9 (right panels) dpf; maximum projection, dorsal view. Nuclei are stained with Hoechst (blue channel). **(E)** *In vivo* quantification from 3 dpf to 9 dpf of the percentage of *mon^NQ039^* mutants (left graph) or WT controls (right graph) with mCherry+ macrophages in different experimental conditions aiming at removing primitive macrophages: injection of Pu1 Mo, Irf8 Mo, or Clodronate liposomes (Clod. Lip.), and Metronidazole (Mtz)-mediated ablation in the *Tg(mpeg1:Gal4;UAS:NfsB-mCherry)* line; n=21 mutant and 88 WT larvae. **(F-H)** Lateral view of the **(F)** tail region or **(G-H)** thymus region of WT control (top) and *mon* mutants (bottom) expressing the [gata2b:Gal4;UAS:LifeAct-eGFP] (hemogenic/HSPC-derived cells, green channel) and mpeg1:mCherry-F (macrophages, red channel) transgenes, at 4 dpf **(F)**, 4.5 dpf **(G)** and 6.5 dpf **(H)**, max. proj. Arrowheads point at double-positive macrophages in controls (purple) and mutants (white); white stars mark pigment cells. Sib, sibling; mut, mutant; hb, hindbrain; mb, midbrain; np, neuropil; CA, caudal artery; CHT, caudal hematopoietic tissue; CV, caudal vein; T, thymus.

We then used a different approach, in which we did not manipulate primitive macrophage production or survival. We took advantage of the *Tg(gata2b:Gal4; UAS:LifeAct-eGFP)* line, in which eGFP specifically highlights aorta-derived HSPCs over the next few days, albeit somewhat mosaically (Butko et al., 2015). We crossed it with *Tg(mpeg1:mCherry-F)* and monitored in the resulting embryos the first appearance of double positive (GFP+, mCherry-F+) cells, which would represent the first definitive (HSPC-derived) macrophages. At 4 dpf (**Fig. 3F**, upper panel) - and as early as 3.5 dpf (**Fig. S3C**), definitive (mCherry+,GFP+) macrophages were detected in the CHT of WT larvae, whereas *mon* mutants were devoid of any macrophages at this stage (**Fig. 3F**, lower panel). At 4.5 dpf, single- and double-positive macrophages could be spotted in various tissues of WT larvae, including the thymic region (**Fig. 3G**, upper panel); macrophages were still absent from *mon* mutants at this stage (**Fig. 3G**, lower panel). Finally, starting at 6.5 dpf, numerous macrophages were detected in the thymic and kidney regions of *mon* mutants, and many of them were double-labelled, evidencing their definitive (HSPC-derived) origin (**Fig. 3H**, lower panel, and Movie 1). So, whether or not we experimentally perturbed primitive macrophage production or survival, in either case the first definitive (HSPC-derived) macrophages arose by 3.5-4 dpf in WT, and only by 6-6.5 dpf in Trim33-deficient mutants.

### Trim33 deficiency affects primitive macrophage survival and definitive macrophage production cell-autonomously

We have thus far established that in Trim33-deficient mutants, primitive macrophages die prematurely - between 3 and 5 dpf and that, independently, the production of definitive macrophages is delayed by at least 2 days.

To assess whether these effects of Trim33 deficiency on macrophages are cell-autonomous, we performed an embryonic parabiosis experiment (Demy et al., 2013), in which we fused WT *Tg(mpeg1:eGFP-F)* blastulas with *mon^TB222^ Tg(gata1:DsRed; mfap4:turquoise)* blastulas (the gata1:DsRed transgene allowed us to sort whether or not the parabiote obtained from *mon^tb222^* heterozygous carrier parents was a *moonshine* homozygous mutant, i.e. producing no circulating Dsred+ primitive erythrocytes). Primitive macrophages originating from both parabiotes/genetic backgrounds were able to colonize, survive and coexist in both parabiotes in WT/sibling (**Fig. 4A**) as well as in WT/mutant embryo combinations (**Fig. 4B**). We monitored mutant *mon^TB222^* /mfap4:turquoise+ macrophages in the WT embryo (**Fig. 4C**, left panel), and WT mpeg1:eGFP-F+ macrophages in the mutant embryo (**Fig. 4C**, right panel) over time. At 2 dpf, both macrophages populations had been able to transcolonize and settle in the other parabiote. By 4 and 5 dpf, WT macrophages had survived and greatly expanded in the mutant parabiote, whereas mutant macrophages had gradually decayed in the WT parabiote. These observations demonstrate a cell-autonomous role for Trim33 in both the survival of primitive macrophages, and the onset of definitive macrophage production.

**Fig. 4.**
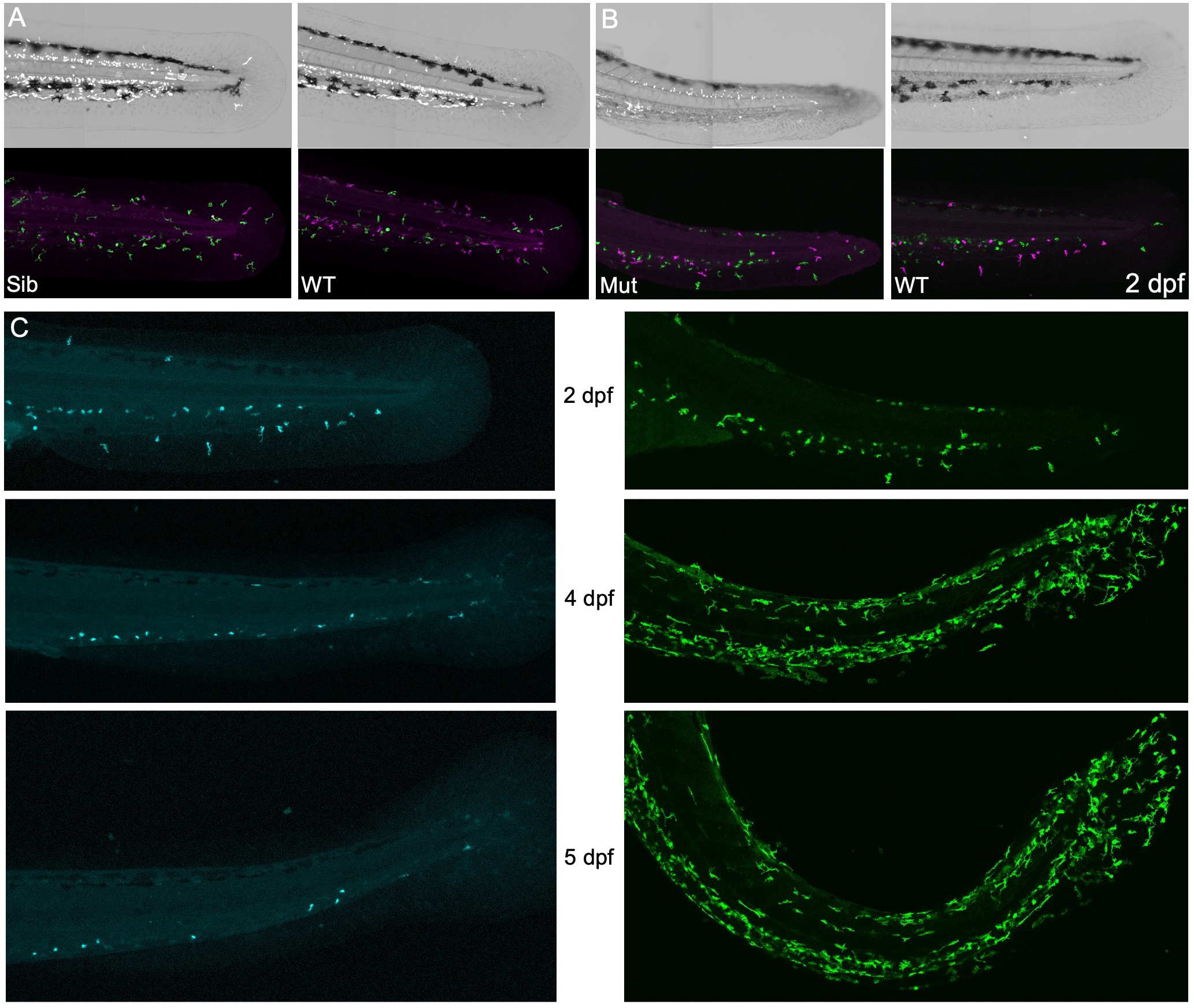
Trim33 deficiency affects the survival of primitive macrophages and production of definitive macrophages cell-autonomously. **(A-C)** Lateral view of the caudal regions of parabiotes fused by the head. **(A)** WT *Tg(mpeg1:eGFP-F)* control embryo (right) fused to a *mon^TB222^/Tg(gata1:Dsred;mfap4:turquoise)* sibling embryo (left) at 48 hpf; **(B)** WT *Tg(mpeg1:eGFP-F)* control embryo (right) fused to a *mon^TB222^/Tg(gata1:Dsred; mfap4:turquoise)* mutant embryo (left). Top images show Gata1:DsRed fluorescence in white, on a bright-field background; bottom images show mpeg1:GFP in green, and mfap4:turquoise in magenta. **(C)** Mutant mfap4:turquoise+ macrophages in the WT tail of the parabiosis (left panels) and WT mpeg1:eGFP-F+ macrophages in the *mon^TB222^* mutant tail of the parabiosis (right panels) at 2, 4 and 5 dpf. Sib, sibling; Mut, mutant

### Definitive granulopoiesis and thrombopoiesis are boosted in Trim33 mutants

In *moonshine* mutants, we further found that the delay in definitive macrophage production correlates with a higher level of definitive neutrophil production, as documented *in vivo* (**Fig. 5A,B**). Whole-mount Sudan Black staining confirmed that these overproduced neutrophils differentiate properly, as they develop fully-stained mature granules, including in the kidney marrow niche (**Fig. 5C**).

**Fig. 5.**
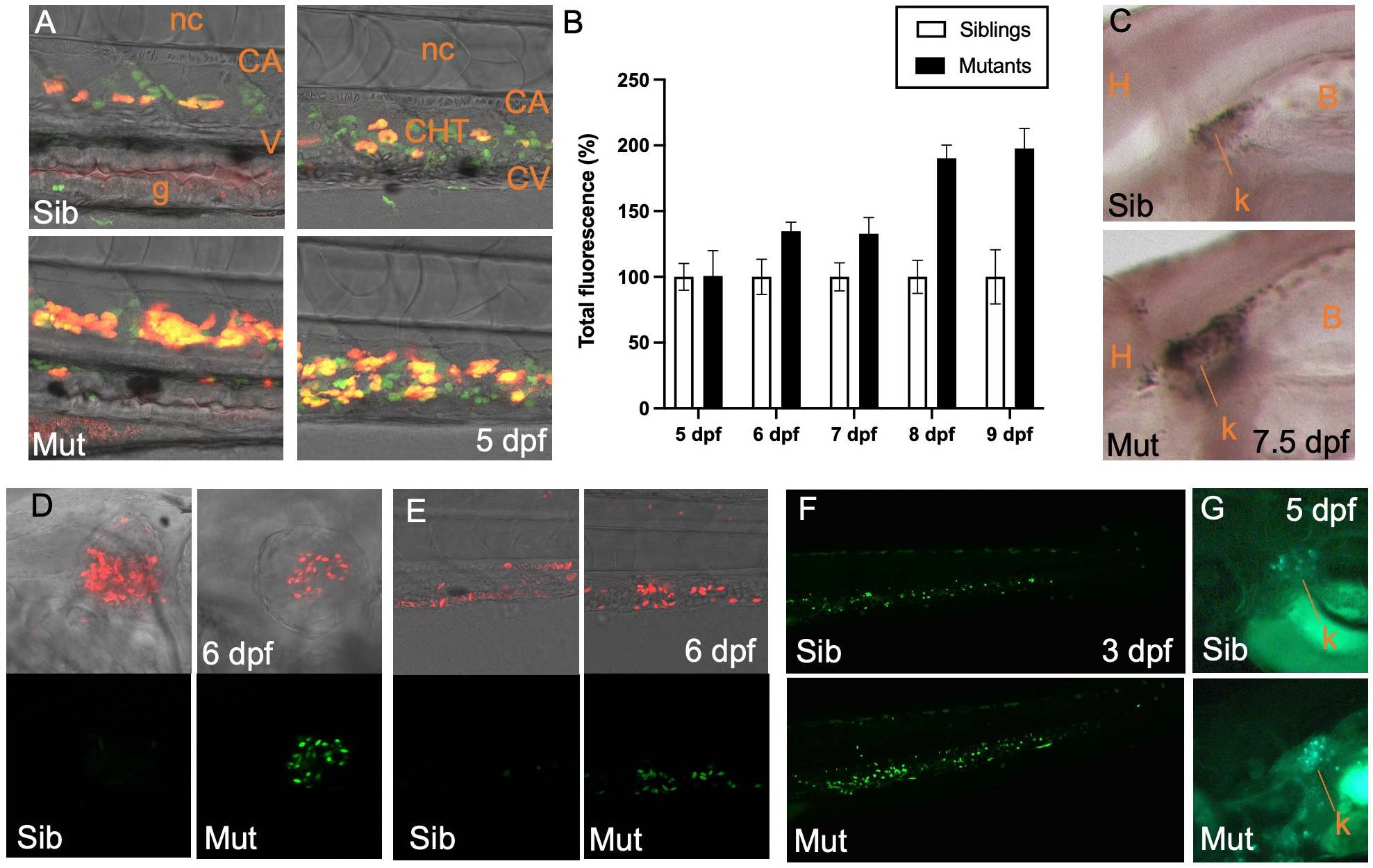
Definitive granulopoiesis and thrombopoiesis are boosted in Trim33-deficient mutants. **(A)** Lateral view of the trunk (left panel) and CHT (right panel) at 5 dpf of control siblings (top) and *mon^NQ039^* mutants (bottom) expressing the coro1a:eGFP (leucocytes, green channel) and lysC:DsRed (neutrophils, red channel) transgenes. **(B)** Quantification of neutrophil-associated fluorescence in live *mon^TB222^/Tg(mpx:eGFP)* siblings and mutants from 5 to 9 dpf, based on total fluorescence (sum of pixel intensities per larva); n=33 mutant and 33 WT larvae. **(C)** Whole-mount Sudan Black staining of mature neutrophils in *mon^NQ039^* siblings (top) and mutants (bottom) at 7.5 dpf; lateral view of the kidney area – rostral to the left. **(D,E)** Lateral view of the cardiac (D) and caudal region (E) of 6 dpf live control sibling (left) and mutant (right) larvae expressing the gata1:DsRed (erythrocytes and thrombocytes, red channel) and CD41:eGFP (HSPCs and thrombocytes, green channel) transgenes, pre-treated with BDM to stop the heartbeat prior to imaging. **(F)** Caudal region of 3 dpf zebrafish *Tg(CD41:eGFP)* larvae control sibling and mutant larvae. **(G)** Kidney region of 5 dpf *Tg(CD41:eGFP)* larvae control sibling and mutant larvae. Sib, sibling; mut, mutant; k, kidney; H, head; b, swim bladder; h, heart; DA, dorsal aorta; CA: caudal artery; CHT, caudal hematopoietic tissue; CV, caudal vein; nc, notochord; g, gut

In addition, from 3 dpf onwards, *moonshine* larvae recovered gata1:Dsred+ circulating cells, but it appeared that these cells were mostly not erythrocytes, but thrombocytes, as they were also CD41:eGFP+ (**Fig. 5D,E**). *moonshine* mutants produced a lot more thrombocytes than their siblings (**Fig. 5D,E** and **Movie 2**), from both the CHT and kidney marrow (**Fig. 5F,G**). These new observations suggest a disruption in the balance of HSPC-derived macrophage vs. neutrophils and erythrocytes vs. thrombocytes production in *moonshine* mutants, thus uncovering a new perspective regarding the central role of Trim33 as a hematopoietic regulator.

## DISCUSSION

In this study we have found that in Trim33-deficient zebrafish, macrophages of the primitive lineage disappear prematurely, and that the production of new macrophages from the definitive hematopoiesis (DH) is delayed. The combination of these two features leads to a complete depletion of macrophages throughout the swimming larva by 4 and 5 dpf. Recent lineage tracing studies have shown that primitive microglia, derived from primitive macrophages, still predominates in the zebrafish brain by 25 dpf, and is then progressively replaced by DH-derived microglia over the following weeks (Ferrero et al., 2018; Xu et al., 2015). Similarly, the primitive macrophages that settle in the epidermis still predominate by 6 dpf, and are progressively replaced by DH-derived macrophages over the next week (He et al., 2018). Here we have found that interstitial macrophages wandering in the mesenchyme are still present by 11 dpf. We previously showed that in *moonshine* embryos, primitive macrophages and neutrophils disperse through the embryo, but are unable to navigate according to developmental or inflammatory signals. Notably, the macrophages do not colonize the brain and retina, a chemokine-driven process (Herbomel et al., 2001; Wu et al., 2018). Thus both macrophages and neutrophils essentially remain in the interstitium between organs/tissues (Demy et al., 2017). In contrast, the later demise of primitive macrophages that we document in the present study does not apply to neutrophils. Early signs of this demise appeared to be a less elongated and ramified morphology, and a loss of expression of macrophage-specific genes – the endogenous *mcsfr1* gene, and the *mpeg1* and *mfap4* promoter driven reporter transgenes. Indeed, the fluorescence of farnesylated (hence membrane-targeted) mCherry expressed from the mpeg1 or mfap4 promoter faded rapidly, first disappearing from the plasma membrane while it was still detected in intracytoplasmic membrane compartments (where the mCherry-F signal is highest, both in WT and mutant macrophages). A the same time, the turquoise fluorescent reporter, expressed throughout the cell from the same promoter, was still detected uniformly within the mutant macrophages; that is why it is actually this reporter that allowed us to document their subsequent apoptotic death via time-lapse *in vivo* imaging. We suspect that the membrane-bound mCherry-F protein has a shorter half-life in macrophages due to the huge membrane turnover that characterizes these cells. Even though the mCherry-F signal is highest in intracellular membrane compartments in macrophages of both mutant and WT larvae, these compartments seem to have distinctive features in the mutant macrophages; they appear to mostly coincide with numerous highly refringent and acidic particles – suggesting they could possibly result from their earlier extensive efferocytosis of the dead primitive erythroid progenitors. We know indeed that prior to their demise, the Trim33-deficient primitive macrophages were phagocytically competent and fully handled the elimination of the primitive erythroid progenitors, that underwent apoptosis during somitogenesis (Demy et al., 2017, and data not shown). Yet this possibly challenging task early in macrophage development cannot be per se the cause of their later premature death. Indeed in our parabiosis experiment in which a *moonshine* embryo is fused with a WT sibling, both the WT and mutant primitive macrophages were exposed to the apoptotic bodies from the mutant primitive erythroid progenitors; yet only the mutant macrophages gradually decayed between 2 and 5 dpf. This demonstrates that Trim33 is required cell-autonomously for the lifespan of the yolk sac derived primitive macrophages in zebrafish. In mammals, yolk sac derived macrophages have been shown to constitute the majority of resident macrophages in many adult tissues throughout life (Kierdorf et al., 2015). Our results suggest that Trim33 could well be one of the key transcriptional regulators fostering their remarkably long lifespan and self-renewal capacity.

We then found that in *moonshine* mutants, the first production of definitive macrophages (from aorta-derived HSPCs) is delayed by at least 2 days relative to WT. To establish that, we first had to determine when the first definitive macrophages normally arise, as this was not known. One reason is that the first macrophages (and neutrophils) to be found in the CHT niche are actually of primitive origin. Indeed when the primitive blood circulation starts by 25 hpf, in the yolk sac it is not yet enclosed in a vessel; hence it bathes the primitive myeloid cells that are still maturing there, (Herbomel and Levraud, 2005; Herbomel et al., 1999); part of them are taken by the blood flow, and most of these home in the tail, within the forming CHT vasculature (Le Guyader et al., 2008; Murayama et al., 2006). Therefore to discern the beginning of definitive macrophage and neutrophil production in the CHT, it was needed either to first get rid of the interference of the primitive myeloid cells, or to use a cell labelling method able to discriminate the two origins. We did both approaches, and they led to the very same results – the first definitive monocytes/macrophages arise in the CHT by 3.5 dpf. This consistency notably tells us that the absence of primitive macrophages did not trigger e.g. some acceleration of macrophage production from definitive hematopoiesis. As aorta-derived HSPCs begin to home in the CHT by 1.5 dpf (Kissa et al., 2008), this delay could mean that the differentiation process from freshly born HSPCs to monocytes/macrophages takes a minimum of 2 days (of note, while in mammals macrophage production from definitive hematopoiesis is known to occur through a monocyte intermediate, in the zebrafish model we do not yet have markers to discriminate monocytes from macrophages within hematopoietic organs).

In contrast, in *moonshine* larvae, the first definitive macrophages arose no earlier than 6 dpf, not particularly in the CHT but rather in the anterior part of the larvae, notably the thymus and anterior kidney areas. Since the definitive HSPCs have begun to colonize the final hematopoietic organ, the anterior kidney, by 4 dpf, these observations could suggest that in Trim33-deficient larvae, monocyte/macrophage production from the definitive HSPCs may not occur at all in the CHT, but only later in the kidney marrow.

In addition to the delayed (and possible absence of) production of definitive macrophages in the CHT of Trim33-deficient larvae, we found that their CHT overproduces not only neutrophils, but also thrombocytes, relative to WT larvae. Based on the defective erythropoiesis and enhanced granulopoiesis in that niche, Monteiro et al. (Monteiro et al., 2011) proposed that Trim33 deficiency caused an imbalance between the myeloid and erythroid fates for the HSPCs in the CHT. The more complete view brought up by our results, i.e. the concomitant lack of macrophage production and excess of thrombocytes, may suggest an alternative hypothesis - an imbalance between the erythroid and thrombocytic fates on one side, and between the macrophage and neutrophil fates on the other side. However, the impact of Trim33 deficiency on HSPC differentiation output is still different in the final niche, the kidney marrow, where erythropoiesis is still compromised (Ransom et al. 2004), neutrophils and thrombocytes are still overproduced, but monocytes/macrophages are now produced as well. It will be interesting to learn whether a differential impact of Trim33 deficiency in the two niches also occurs in the homologous hematopoietic niches of mammals – the fetal liver then the bone marrow.

## MATERIALS AND METHODS

### Zebrafish lines and embryos

Wild-type, transgenic and mutant zebrafish embryos were raised at 28°C in Embryo Water (Volvic^©^ water containing 0.28 mg/mL Methylene Blue [M-4159; Sigma] and 0.03 mg/mL 1-phenyl-2-thiourea [P-7629; Sigma]), and staged according to Westerfield (Westerfield, 2007). The *moonshine^NQ039 or t30813^* (Demy et al., 2017) and *moonshine^TB222^* (Ransom et al., 2004) mutations were maintained by outcross of heterozygous carriers with WT and/or Tg fish from the following lines: Tübingen wildtype (WT) fish (ZIRC), *Tg(mpeg1:mCherry-F)^ump2^* (Nguyen-Chi et al., 2014), *Tg(lyz:DsRed2)^nz50^* (Hall et al., 2007), *Tg(mfap4:mCherry-F)^ump6^* (Phan et al., 2018), *Tg(mpeg1:Gal4FF)^gl25^* (Palha et al., 2013), *Tg(UAS:Kaede)^rk8^* (Hatta et al., 2006), *Tg(mfap4:turquoise)^xt27^* (Walton et al., 2015), *Tg(UAS-E1b:Eco.NfsB-mCherry)^c264^* (Davison et al., 2007), *Tg(gata2b:Gal4;UAS:lifeAct-eGFP)* (Butko et al., 2015), *Tg(mpeg:GFP-F)^sh425^* (Keatinge et al., 2015), *Tg(mpx:GFP)^i114^* (Renshaw et al., 2006), *Tg(gata1:dsRed)^sd2^* (Traver et al., 2003), and *Tg(CD41:GFP)^la2^* (Lin et al., 2005).

### Identification of the *moonshine^NQ039^* mutation of the Trim33 gene

Pools of ~100 zebrafish larvae from 4 replicates *moonshine^NQ039^* couples were anesthetized with 0.16 mg/ml Tricaine [A-5040; Sigma], screened for the *mon^NQ039^* phenotype (based on absence of circulating erythrocytes) and their total RNA was extracted using TRIzol (Invitrogen), following the manufacturer’s protocol. cDNA was obtained using MMLV H Minus Reverse Transcriptase (Promega) with a dT17 primer. Trim33 PCR was performed using LA Taq polymerase (Takara) and the following primer pair:

5’-GCTCCTACTGCTCCTCCGTCCA-3’
5’-GATGGACGAGCTGGAGTGTG-3’

Sequencing of the amplicon was performed by Eurofins using the forward primer, and led to the identification of a C>T change (Gln>Stop) at the 1745 bp position, resulting in a truncated protein retaining 581 out of the 1176 amino acids of the WT protein. The other *moonshine* mutant used in this study,*mon^tb222^* (Ransom et al., 2004), harbors a nonsense mutation resulting in a Trim33 protein retaining the first 433 amino acids.

### FACS

Zebrafish *Tg(mpeg1:mCherry-F)* larvae obtained from *mon^NQ039^* heterozygous parents were anesthetized at 4 dpf with 0.16 mg/ml Tricaine [A-5040; Sigma], screened for the *mon^NQ039^* phenotype (based on absence of circulating erythrocytes) then dissociated into single-cell suspensions as previously described (Covassin et al., 2006). Fluorescence-activated cell sorting (FACS) of red fluorescent macrophages was performed using a FACSCALIBUR and a FACSAria cytometers (BD Biosciences). Data analysis was performed using FlowJo software (TreeStar). The experiment was replicated 3 times, using pools of >50 larvae per condition.

### Whole-mount mRNA in situ hybridization and immunohistochemistry

Embryos and larvae were anesthetized with Tricaine at the stage of interest, then fixed overnight at 4°C in 4% methanol-free formaldehyde (Polysciences, # 040181). Wholemount *in situ* hybridization (WISH) was performed according to Thisse and Thisse, 2014 (Thisse and Thisse, 2014). Whole-mount immunohistochemistry (WIHC) was performed as described previously (Murayama et al., 2006), omitting the acetone treatment. The primary antibody used was a rabbit anti-zebrafish L-plastin polyclonal antibody (at 1:5000) (Le Guyader et al., 2008) and the secondary antibody a Cy3-coupled anti-rabbit antibody (111-166-003; Jackson Immunoresearch), used at 1:800 dilution.

### Sudan Black staining of neutrophils

Sudan Black staining of neutrophils in formaldehyde fixed embryos was done as previously described (Le Guyader et al., 2008).

### UV photoconversion

Embryos from the *Tg(mpeg1:gal4; UAS:Kaede)* line were dechorionated at 2 dpf, then screened under a Leica M165 FC fluorescence stereomicroscope. Embryos were exposed in groups of 10 under UV light to photoconvert Kaede from green to red using a HBO lamp, and their macrophages were then followed up every day from 2 to 11 dpf. All larvae were kept in the dark and observed with a UV filter during the whole duration of the study.

### Pharmacological treatments & live staining

Staining of acidic compartments was achieved by bathing live zebrafish embryos for >30min in the dark in a LysoID solution (Enzo, #510005) diluted at 1/1000e. Larvae were then rinsed 3 times, and quickly imaged. For macrophages ablation with Metronidazole (Mtz), embryos were dechorionated and screened for red fluorescence using a fluorescence stereomicroscope. They were incubated in the dark at 28°C overnight with 10mM Mtz freshly diluted from powder (Sigma M3761) in embryo water + 1% DMSO. Embryos were washed 3 times before observation/imaging. To arrest the heartbeat so as to image the heart content, 6 dpf zebrafish larvae were placed for a few hours in water containing 10 mM BDM (2.3-butanedione monoxime: Sigma, B-0753) as previously described (Miyasaka et al., 2011).

### Morpholino injections

The two antisense morpholinos (Mos) used in this study were synthesized by Gene Tools:

Anti-Pu.1 MO (Rhodes et al., 2005): 5’-GATATACTGATACTCCATTGGTGGT-3’
Anti-Irf8 MO (Yu et al., 2017): 5’-AATGTTTCGCTTACTTTGAAAATGG-3

1-5 nL of 0.6 mM Mo solution was microinjected in one-to two-cell stage embryos.

### Clodronate liposomes intravenous injection

Clodronate liposomes (clodronateliposomes.com) were stained with a DiO solution (2.5 mg DiO in 1mL ethanol). 10 μL of DiO solution were added per mL of liposomes solution. The sample was incubated for 10 min at room temperature, then centrifuged twice at 20.000 g for 10 min. The supernatant was removed and replaced with sterile PBS. A few nanoliters of the final solution was injected in the Duct of Cuvier of 48 hpf zebrafish embryos, and proper injection in the blood flow was subsequently verified under a fluorescence dissecting scope.

### Parabiosis

Parabiotic embryos were generated as previously described (Demy et al., 2013) by fusing WT *Tg(mpeg1:GFP-F)* embryos at the late blastula stage to stage-matched *mon^NQ039^ Tg(gata1a:DsRed; mfap4:turquoise)* embryos; mutant embryos were identified by the absence of circulating DsRed-positive erythroid cells.

### Microscopy and image analysis

Low-magnification bright-field images were acquired using video cameras mounted on a Leica Macrofluo driven by the Metavue (Metamorph) software or a Zeiss Macrofluo driven by the Zen software (Zeiss). Wide-field video-enhanced (VE) Nomarski / differential interference contrast (DIC) and fluorescence microscopy were performed as described previously (Herbomel and Levraud, 2005; Murayama et al., 2006), through the 40x/1.00 water-immersion objective of a Nikon 90i microscope or the 40x/1.00 oil-immersion objective of a Reichert Polyvar 2 microscope. Images were obtained from a HV-D20 3-CCD camera (Hitachi), digitized through a GVD-1000 DV tape recorder (Sony), then still images were collected using the BTVpro software (Bensoftware, London). For fluorescence confocal imaging, embryos and larvae were mounted as previously described (Demy et al., 2017). Images were then captured at selected times on an inverted Leica SP8 set-up allowing multiple point acquisition, so as to image mutants and their siblings in parallel. Images were treated and analyzed using the Fiji (ImageJ) software.

### Macrophages morphological analysis & cell counting

#### Macrophage count

Cells were manually counted on Fiji (ImageJ) from maximum projections of total z-stacks of whole live zebrafish larvae, obtained with the SP8 Leica confocal set-up.

#### Larvae with/without macrophages

At the stages of interest, live mutant & WT larvae were observed under a Leica M165FC fluorescence stereomicroscope and binary scored as having a “normal” macrophage population or not, according to the phenotype presented in Fig. 1A.

<u>Fluorescence intensity</u> was automatically measured on Fiji (ImageJ) after manual thresholding of images obtained with the SP8 Leica confocal set-up. It was then expressed in Area (number of pixels above threshold) and Pixel Intensity.

<u>Macrophage roundnes</u>s was automatically measured on Fiji (ImageJ) from images obtained with the SP8 Leica confocal set-up, after manual determination of cells.

<u>Macrophage ramifications</u> were manually counted on Fiji (ImageJ) from maximum projections of Z stacks of live zebrafish larvae, obtained with the SP8 Leica confocal set-up.

### Statistical analysis

To evaluate differences between means of non-Gaussian data, they were analyzed with a Mann-Whitney-Wilcoxon. When appropriate, distributions were normalized by log, and an analysis of variance (Two-way ANOVA) was performed (for experiments with two variables). *P*<0.05 was considered statistically significant (symbols: ****, P<0.0001; ***, *P*<0.001; **, *P*<0.01; *, *P*<0.05). Statistical analyses and graphic representations were done using Prism software.

## Acknowledgements

We wish to thank our fish facility team for their excellent care of the fish, Léa and Valerio Laghi for their help in distant work procedures.

## Author contributions

Conceptualization: D.L.D., M.T., P.H.; Methodology: D.L.D., P.H.; Validation: D.L.D., M.T., I.L., W.G., P.H.; Formal analysis: D.L.D., P.H.; Investigation: D.L.D., A-L.T., M.L., M.T., L.C., C.P.; Resources: D.L.D., M.T., P.H.; Data curation: D.L.D., P.H.; Writing - original draft: D.L.D., P.H.; Writing - review & editing: D.L.D., P.H.; Visualization: D.L.D., A-L.T., M.L., M.T., P.H.; Supervision: P.H.; Project administration: P.H.; Funding acquisition: P.H.

